# Molecular mechanisms underlying the early steps of floral initiation in seasonal flowering genotypes of cultivated strawberry

**DOI:** 10.1101/2025.01.20.633581

**Authors:** Freya MR Ziegler, Amèlia Gaston, Karine Guy, Marie Devers, Erika Krüger, Bastienne Brauksiepe, Klaus Eimert, Sonia Osorio, Béatrice Denoyes, Björn Usadel

## Abstract

Floral initiation is required for sexual reproduction in angiosperm plants, and has a significant impact on crop yields. In cultivated strawberry, the molecular basis of floral initiation is poorly understood and most studies have focused on a single genotype grown under controlled conditions. To gain more insight into this process, we conducted a study under natural conditions in two countries using two seasonal flowering cultivars. We focused on the early steps of floral initiation by using samples spanning key developmental stages of the shoot apical meristem. The analysis of differential gene expression in leaf and terminal bud tissues revealed enrichment for genes involved in carbohydrate metabolism and phytohormone pathways in leaves. Other protein classes that were enriched during early floral initiation were associated with cytoskeleton organization, cell cycle regulation, and chromatin structure. We also identified genes associated with the photoperiodic pathway, well-characterized floral integrators such as TFL1 and SOC1, and several linked to phytohormone regulation, such as *XTH23*, *PP2* and *EIN3*.

## Introduction

Floral initiation is a pivotal event in the life cycle of angiosperm plants that marks the transition from vegetative to reproductive development. It occurs in the shoot apical meristem (SAM), which generates leaves, shoots and flowers (Benlloch et al., 2007; Figure 1A). Flowering is necessary for reproductive success, significantly affecting pollination, fruit production, and crop yields (Weberling 1989; Wyatt, 1982).

**Figure 1:**
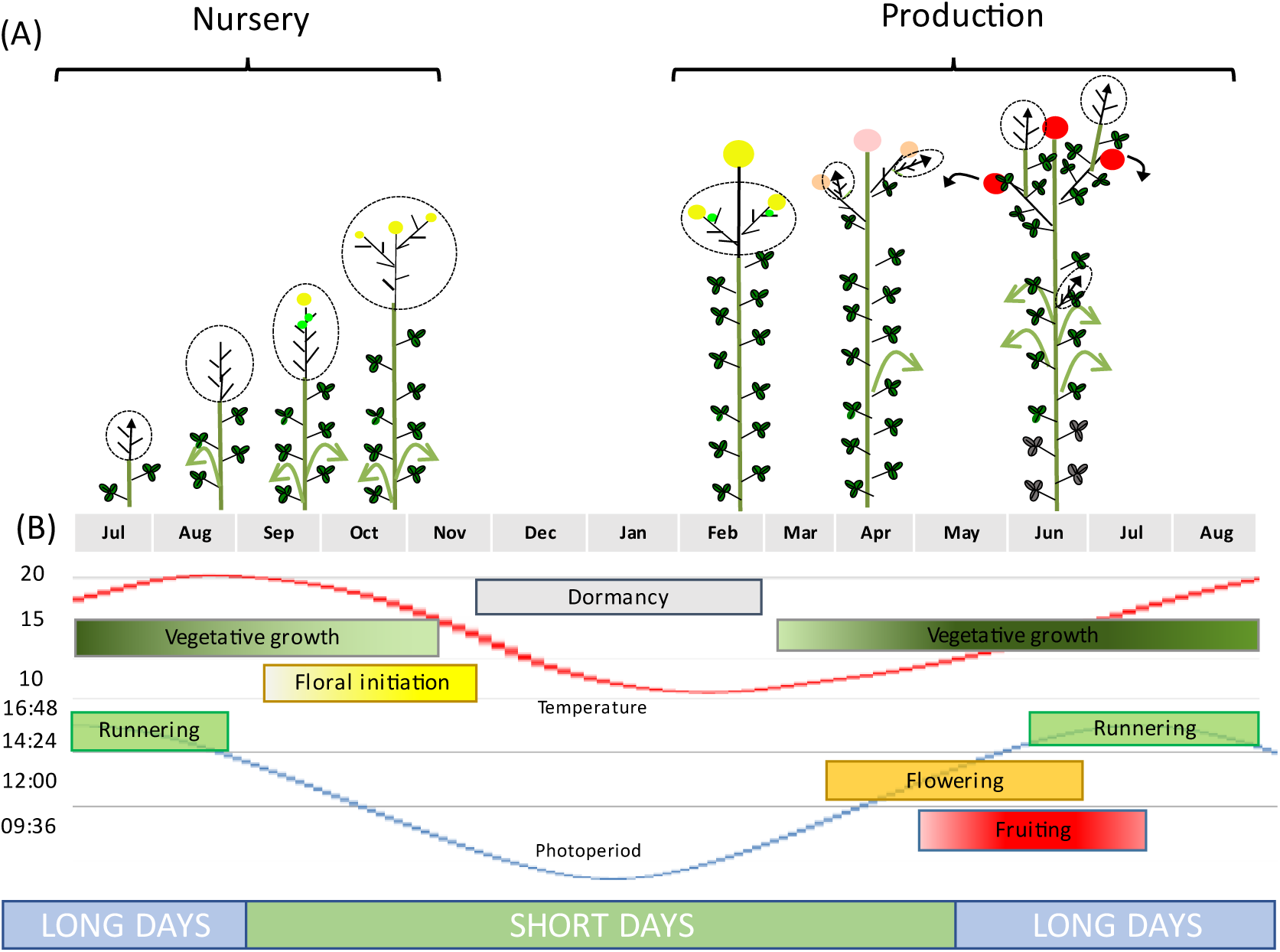
Developmental processes in seasonal flowering varieties of cultivated octoploid strawberry. (A) In the nursery, temperatures and day length decline, allowing floral initiation to occur. The shoot apical meristem transitions from vegetative (arrow) to floral development (yellow sphere). Because strawberry is a rosette, the terminal bud is not visible and is represented here within the dotted line at the base of the plant. (B) Runnering occurs on long days. Dormancy occurs at the end of autumn and during winter, when temperatures do not allow vegetative development. The dashed circle indicates the terminal bud, which includes the foliar primordia.

In strawberry (*Fragaria × ananassa*), the most widely cultivated berry crop, the fruit yield depends on the timing and duration of floral initiation (Costes et al., 2014). Most strawberry genotypes are seasonal flowering (also called short-day, single cropping or June bearing) varieties, with a single fruiting period in the spring of year *N*. Floral initiation occurs during the previous autumn (year *N* – 1) and is triggered by declining temperatures and day length at the end of summer/beginning of autumn (Heide et al., 2013; Figure 1B). Floral initiation is followed by floral development, specifically the organogenesis of flowers in the inflorescence. Following dormancy and the fulfilment of chilling requirements, flowers initiated in autumn begin to emerge in spring.

Strawberry plants also reproduce asexually via stolons, which are elongated stems bearing daughter plants (Tenreira et al., 2017). The choice between sexual and asexual reproduction is made in the axillary meristem, which can produce either an inflorescence-bearing branch or a stolon, and thus determines two antagonistic traits: fruit yield, a major trait for producers, and daughter-plant yield, a major trait for nurseries (Tenreira et al., 2017; Gaston et al., 2021).

The interplay between photoperiod and temperature, and their combined influence on floral initiation have been extensively documented (Stewart and Folta 2010; Heide et al. 2013). The overarching consensus is that optimal conditions for floral initiation involve temperatures of 12–18°C and photoperiods of 10–12 h maintained over a period of 3–4 weeks. Global radiation also contributes to floral initiation (Krüger et al., 2022). Variations in these parameters are mainly due to cultivar-specific responses (Verheul et al., 2006; Krüger et al., 2022).

Recent work has begun to identify and characterize the strawberry genes controlling floral initiation, plant architecture and yield (Hytonen and Kurokura, 2020). The CENTRORADIALIS/TERMINAL FLOWER 1/SELF-PRUNING (CETS) family, represented by *TFL1/FT* genes, plays a pivotal role (Wickland and Hanzawa, 2015; Kurokura et al., 2017; Gaston et al., 2021). TFL1 acts as a floral repressor in both diploid *Fragaria vesca* (Iwata et al., 2012; Koskela et al., 2012) and cultivated strawberry (Koskela et al., 2016), and its expression declines during floral initiation in cultivar Benihoppe (Liang et al., 2022). Three strawberry *FT* genes have also been identified. *FveFT1* encodes a long-day floral activator, as shown in the *tfl1* diploid genetic background (Koskela et al., 2012; Rantanen et al., 2014). *FveFT2* encodes a non-photoperiodic florigen (Gaston et al., 2021) and operates in tandem with the photoperiodic anti-florigen *FveTFL1* (Gaston et al., 2021). Overexpression of *FveFT2* confers a very early flowering phenotype. In octoploid cultivated strawberry, *FanFT3* encodes a floral repressor (Koembuoy et al., 2020), and *FveFT3* overexpression promotes plant branching in the *tfl1* diploid genetic background.

The regulation of flowering is intricately linked to the production of stolons. Gibberellin biosynthesis and signaling affect strawberry plant architecture and fruit yield by specifying whether the axial meristem produces a stolon or an inflorescence-bearing branch crown (Tenreira et al., 2017; Caruana et al., 2018). The natural mutation in *FveGA20ox4* generates a runnerless phenotype (Tenreira et al., 2017), which can be reversed by mutating *FveRGA1* (*REPRESSOR OF GIBBERELIC ACID1*), encoding a DELLA protein (Caruana et al., 2018). The molecular mechanisms governing the balance between sexual and asexual reproduction in the axial meristem are only beginning to be understood.

The molecular control of floral initiation in the cultivated strawberry *F. × ananassa* has yet to be studied in detail under natural conditions. Here, we examined the early molecular events of floral initiation by transcriptomic analysis, taking into account both the genotype and environment by studying two cultivars – the Italian cultivar Clery (CL) and the French cultivar Gariguette (GA) – at two locations, one in France and one in Germany. We analyzed RNA-Seq data from leaf and terminal bud tissues during the transition from the vegetative phase to the early stages of floral initiation in the SAM (weeks 29, 32, 33 and 35). As anticipated, we found that both cultivars were enriched during the early period of floral initiation for differentially expressed genes (DEGs) encoding proteins involved in flowering, but also those relating to chromatin structure, cytoskeletal organization, serine/threonine phosphatase signaling, cell division, cell wall organization, and RNA biosynthesis.

## Results

### Phenotypic variation of floral initiation according to the cultivar and environment

We investigated floral initiation over time by dissecting the terminal bud in cultivars CL and GA and observing the SAM from the middle of July until the end of October over 3 years (2016–2018) in two different environments in France and Germany. The SAM in the terminal bud was always vegetative on the first sampling date in July regardless of the genotype or location (Figure 2). The SAM transitioned to a floral identity when its apical dome rose above the level of the developing stipules (Krüger et al., 2022).

**Figure 2:**
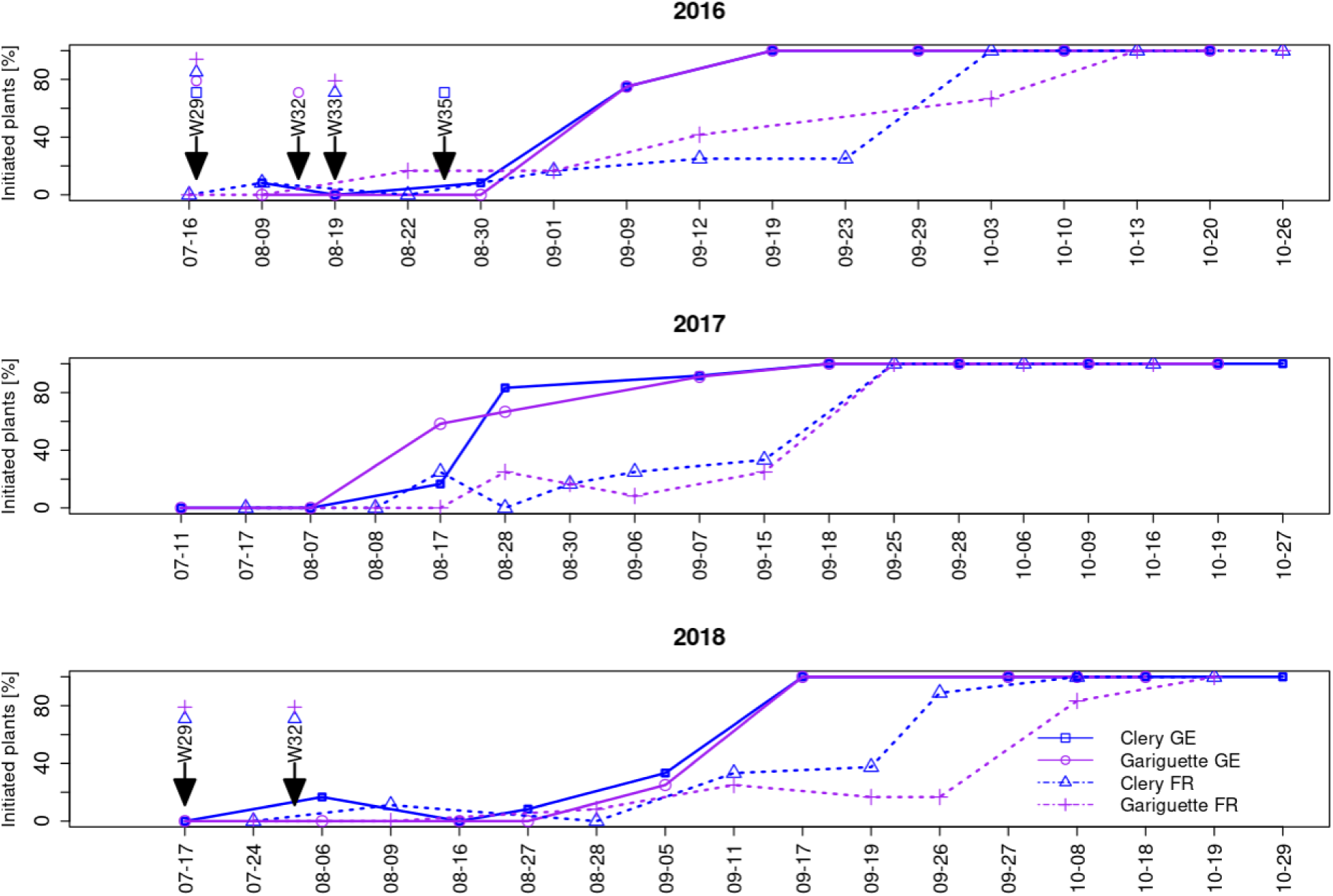
The frequency of plants undergoing floral initiation over time according to cultivar and location. Floral initiation was confirmed when the SAM reached at least stage 2 (terminal bud dome has risen above the developing stipules). Sampling dates for the RNA-Seq analysis of leaf and terminal bud tissues are indicated by arrows labeled with the sampling week: w29 (T0), w32 (T10A), w33 (T10B) and w35 (T50). FR = France, GE = Germany.

Because different plants were tested in each sample, the percentage of initiated plants (those with a floral terminal bud) fluctuated between consecutive sampling dates. The beginning of floral initiation was similar in France and Germany, with a low percentage (<10%) of initiated plants during the first weeks, rising steadily and then jumping from 20–40% to 90–100% in a single week (Figure 2). In Germany, this week was in the middle or at the end of August (depending on the year) whereas in France it was in the middle or at the end of September (Figure 2).

Floral initiation was followed by organogenesis in the SAM, resulting in the formation of an entire inflorescence with differentiated flower organs (still in the terminal bud, Figure 1). SAM development into an inflorescence followed the same tendency every year at both locations, although it occurred earlier for genotype CL than GA, and earlier in Germany than in France, as anticipated given the earlier floral initiation in Germany (Figure 3).

**Figure 3:**
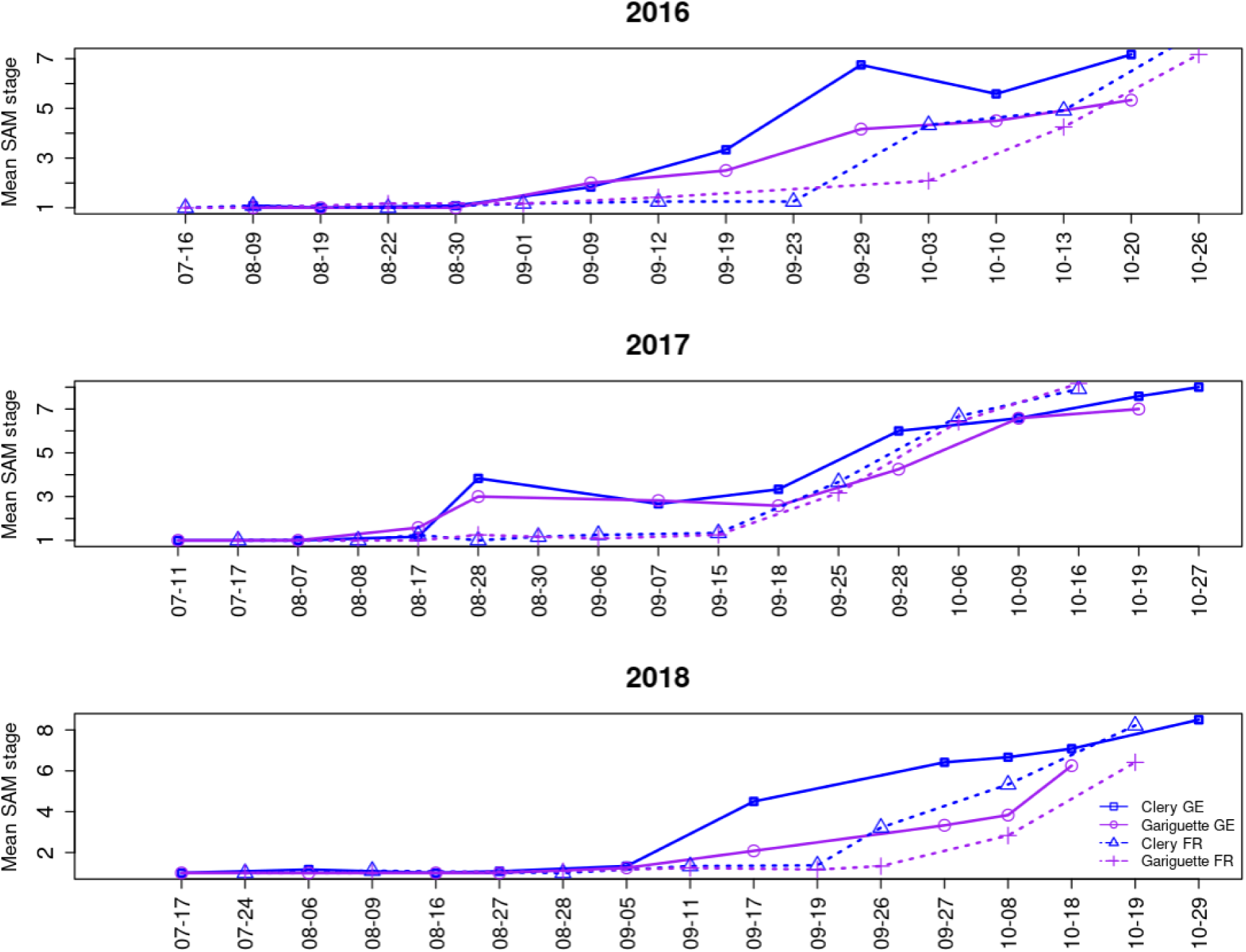
The mean floral developmental stage of the shoot apical meristem over time for the strawberry cultivars Clery and Gariguette in Germany (GE) and France (FR). SAM staging is described in Supplemental Figure S1A.

### Choosing relevant sampling dates to study early floral initiation

We collected samples for RNA-Seq at three time points, when all plants were vegetative: T0 with no plants initiated, T10 when ∼10% of the plants were initiated, and T50 when ∼50% of the plants were initiated. Given the effects of genotype and environment noted above, the time points differed by genotype, location and year (Table 1; Figure 2). For T0 samples, leaves and terminal buds were collected from daughter plants maintained on their mother plants during w29 or w30. The sample names were based on genotype, location/year and sampling time point as follows: CL_GE16_T0, GA_GE16_T0, CL_FR16_T0, GA_FR16_T0, CL_FR18_T0, and GA_FR18_T0.

**Table 1.** Frequency of floral initiation [%] in samples of strawberry leaf (L) and terminal bud (TB) acquired in France (FR) and Germany (GE) over 3 years for the cultivars Clery (CL) and Gariguette (GA).

| ID | Location | Year | Cultivar | Date observation | Tissue | Week | Time point | Floral initiation [%] |
| --- | --- | --- | --- | --- | --- | --- | --- | --- |
| **Starting Point (T0)** |  |  |  |  |  |  |  |  |
| CL_GE16_T0 | GE | 2016 | CL | 2016-07-19 | L, TB | 29 | T0 | 0 |
| GA_GE16_T0 | GE | 2016 | GA | 2016-07-19 | L, TB | 29 | T0 | 0 |
| CL_FR16_T0 | FR | 2016 | CL | 2016-07-16 | L | 29 | T0 | 0 |
| GA_FR16_T0 | FR | 2016 | GA | 2016-07-16 | L | 29 | T0 | 0 |
| CL_FR18_T0 | FR | 2018 | CL | 2018-07-24 | L | 30 | T0 | 0 |
| GA_FR18_T0 | FR | 2018 | GA | 2018-07-24 | L | 30 | T0 | 0 |
| **Initiated plants** |  |  |  |  |  |  |  |  |
| Early floral initiation (~10% of plants initiated) (T10) |  |  |  |  |  |  |  |  |
| CL_GE16_T10 | GE | 2016 | CL | 2016-08-30 | L, TB | 35 | T10 | 8.3 |
| GA_GE16_T10 | GE | 2016 | GA | 2016-08-09 | L, TB | 32 | T10 | 8.3 |
| CL_FR16_T10 | FR | 2016 | CL | 2016-08-22 | L | 33 | T10 | 0 |
| GA_FR16_T10 | FR | 2016 | GA | 2016-08-22 | L | 33 | T10 | 16.6 |
| CL_FR18_T10 | FR | 2018 | CL | 2018-08-09 | L | 32 | T10 | 11.1 |
| GA_FR18_T10 | FR | 2018 | GA | 2018-08-09 | L | 32 | T10 | 0 |

| Floral initiation continued (~50% of plants initiated) (T50) |  |  |  |  |  |  |  |  |
| --- | --- | --- | --- | --- | --- | --- | --- | --- |
| CL_GE16_T50 | GE | 2016 | CL | 2016-09-09 | L | 36 | T50 | 75 |
| GA_GE16_T50 | GE | 2016 | GA | 2016-09-09 | L | 36 | T50 | 75 |
| CL_FR16_T50 | FR | 2016 | CL | 2016-09-12 | L | 37 | T50 | 25 |
| GA_FR16_T50 | FR | 2016 | GA | 2016-09-12 | L | 37 | T50 | 41.1 |

For T10 samples, leaves and terminal buds were collected when the percentage of initiated plants varied from 0% to 17%. A 0% frequency of initiated plants was observed in sample CL_FR16, but the percentage was 8.3% 10 days before, and in sample GA_FR18, where the percentage increased to 11% the week after (Figure 1; Table 1). The sample names were CL_GE16_T10, GA_GE16_T10, CL_FR16_T10, GA_FR16_T10, CL_FR18_T10, and GA_FR18_T10. We recovered four T50 samples: CL_GE16_T50, GA_GE16_T50, CL_FR16_T50, and GA_FR16_T50.

### Transcriptome variation in strawberry tissues during floral initiation

To find differences and similarities between cultivars during early floral initiation, the RNA-Seq samples for leaf and terminal bud were compared using a combination of principal component analysis (PCA) and overrepresentation analysis (ORA). All three replicates of each sample were grouped, confirming their homogeneity. For the leaf samples (Figure 4), the gene expression patterns for cultivars CL and GA grown in France were initially similar (CL_FR16_T0 and GA_FR16_T0) but began to segregate along the PC1 axis by stage T10 (CL_FR16_T10 and GA_FR16_T10). This was attributed to genes associated with photosynthesis, uptake of transition metal ions, phytohormone activity, lipid metabolism, and RNA biosynthesis. Notably, for both cultivars at both locations at stage T10 (w32, w33 and w35), a more pronounced separation was observed along the PC2 axis. The separation patterns of both cultivars aligned closely across these time points (Figure 4).

**Figure 4:**
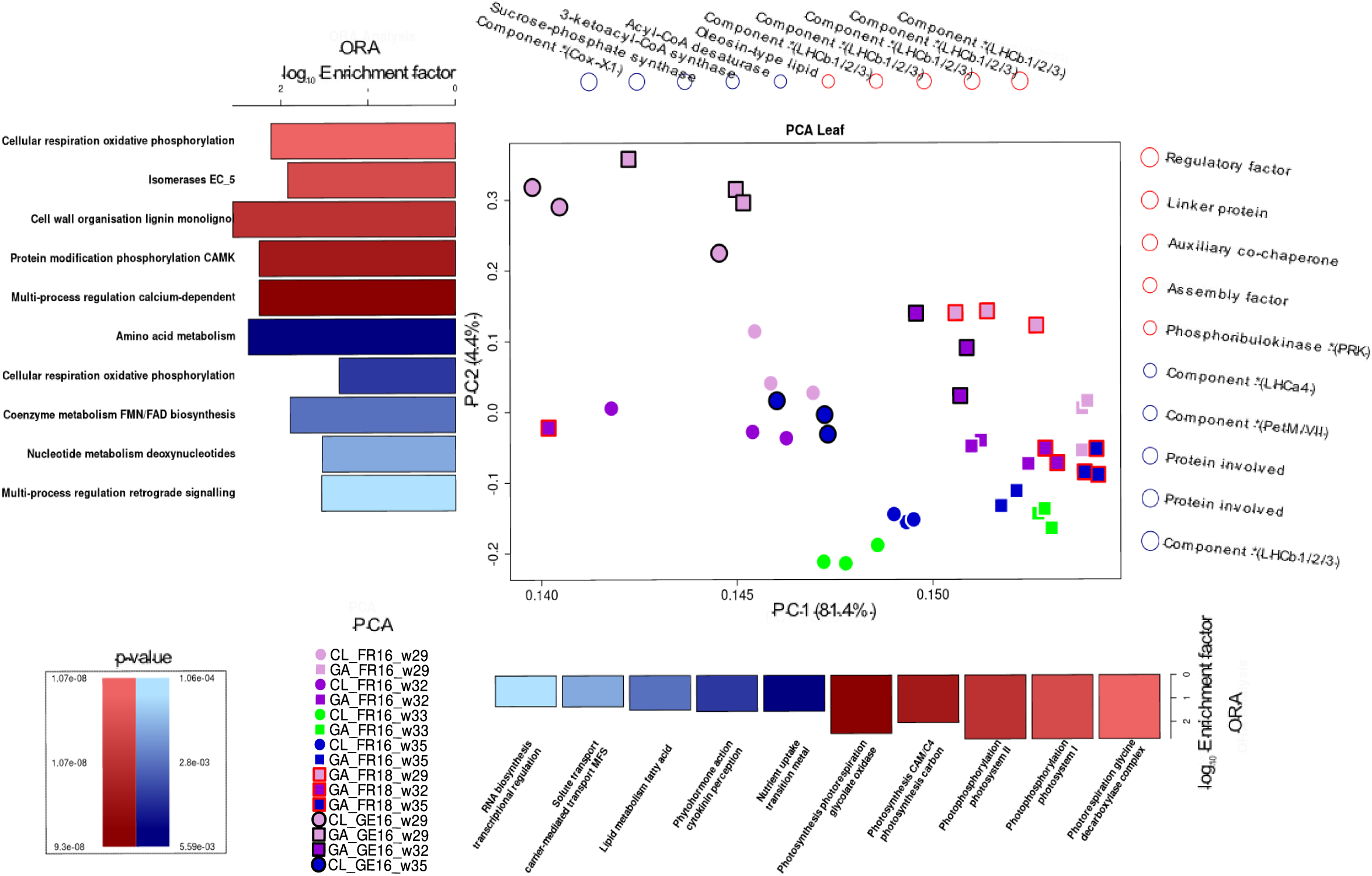
Principal component analysis (PCA) of differentially expressed genes (DEGs) in the leaves of cultivars Clery (CL) and Gariguette (GA) grown in France in 2016 (FR16) and 2018 (FR18), and in Germany in 2016 (GE16), sampled at time points T0 (w29), T10A (w32), T10B (w33) and T50 (w35). Overrepresentation analysis (ORA) was then applied using MapMan protein classes based on the loadings for principal components 1 (PC1) and 2 (PC2). The top 5 MapMan protein classes resulting from ORA (p < 0.01), and their involvement in the separation along PC1 and PC2 in the positive (red) and negative (blue) directions, are shown as bar plots of the log_10_ enrichment factor. Genes strongly associated with the positive (red) or negative (blue) separation determined by PCA are depicted above (PC1) and to the right (PC2). In the PCA plots, circles and squares represent cultivars CL and GA, respectively. Frame colors represent the location/year: none = France 2016, black = Germany 2016, red = France 2018. Block colors represent the stage: light purple = T0, dark purple = T10A, green = T10B, blue = T50.

In Germany, cultivars CL and GA were separated along the PC2 axis from the first time point T0 (CL_GE18_T0 and GA_GE18_T0), particularly for protein classes involved in cellular respiration, cell wall organization, protein modification, and multi-process regulation (Figure 4). But by stage T10, the cultivars were dispersed clusters, which disguised the separation between them (CL_GE18_T10 and GA_GE18_T10A).

For the terminal bud (Supplemental Figure S3), the floral initiation of GA_FR18 was delineated along the PC1 axis. The separation observed at w29, w32 and w35 was mainly influenced by protein classes related to protein homeostasis, biosynthesis, and multi-process regulation of phosphoinositide. Moreover, the segregation at stage T10 (w32) also affected the PC2 axis, and involved protein classes related to cell wall organization (lignin), RNA processing, and secondary metabolism (phenolics). Analogous outcomes were observed for GA_GE16_T0 and GA_GE16_T10, as well as for CL_GE16_T0 and CL_GE16_T50. Although the cultivars were clearly separated along the PC1 axis at stage T0, the results for both cultivars were similar at stages T10 and T50.

### Common DEGs between T0 and T10 during early floral initiation

We focused on early floral initiation under natural conditions by screening for genes that were differentially expressed between the two first time points: T0 (w29) and T10 (w32, w33 or w35 according to the country and the year). We compared pairs of transcript profiles between T0 and T10 samples within each cultivar (CL or GA) and within each organ (leaf or terminal bud). The comparisons are represented by logical names. For example, CL_GE16_T0xT10 refers to the comparison of samples CL_GE16_T0 and CL_GE16_T10. Suffixes L and TB were added to represent leaf and terminal bud tissues, respectively. We identified 71,369 DEGs across all comparisons (FDR-corrected p ≤ 0.05 in all cases).

For leaf samples, the comparison of stages T0 and T10 for cultivar CL grown in Germany (CL_GE16_T0xT10L) and France (CL_FR16_T0xT10L) revealed 7118 DEGs (Supplemental Figure S2A; Supplemental Table S1). The analogous comparisons for cultivar GA (GA_GE16_T0xT10L, GA_FR16_T0xT10L and GA_FR18_T0xT10L) revealed 786 DEGs (Supplemental Figure S2B; Supplemental Table S2). The two cultivars shared 416 common DEGs expressed in leaves (Supplemental Table S3).

For terminal bud samples, the comparisons for cultivar CL (CL_GE16_T0xT10TB) revealed 3385 DEGs (Supplemental Table S4), whereas those for cultivar GA (GA_GE16_T0xT10TB and GA_FR18_T0xT10TB) revealed 1548 DEGs (Supplemental Table S5). The two cultivars shared 1207 common DEGs expressed in the terminal bud (Supplemental Figure S2C; Supplemental Table S6).

Enrichment analysis for MapMan protein classifications was applied to the common DEGs. In CL leaves, the main enriched classes (Figure 5) were plant reproduction, modulation of flowering, enzyme activity, secondary metabolism (terpenoids), and carbohydrate metabolism (starch). Conversely, the main enriched classes in GA leaves were multi-stress responses, cytoskeletal organization (microtubule network), chromatin organization/structure, phytohormone activity, carbohydrate metabolism and RNA biosynthesis (Figure 5). The DEGs shared by both cultivars were enriched for proteins related to solute transport channels, cell wall organization (pectin, rhamnogalacturonan), cell division and photosynthesis (Figure 5).

**Figure 5.**
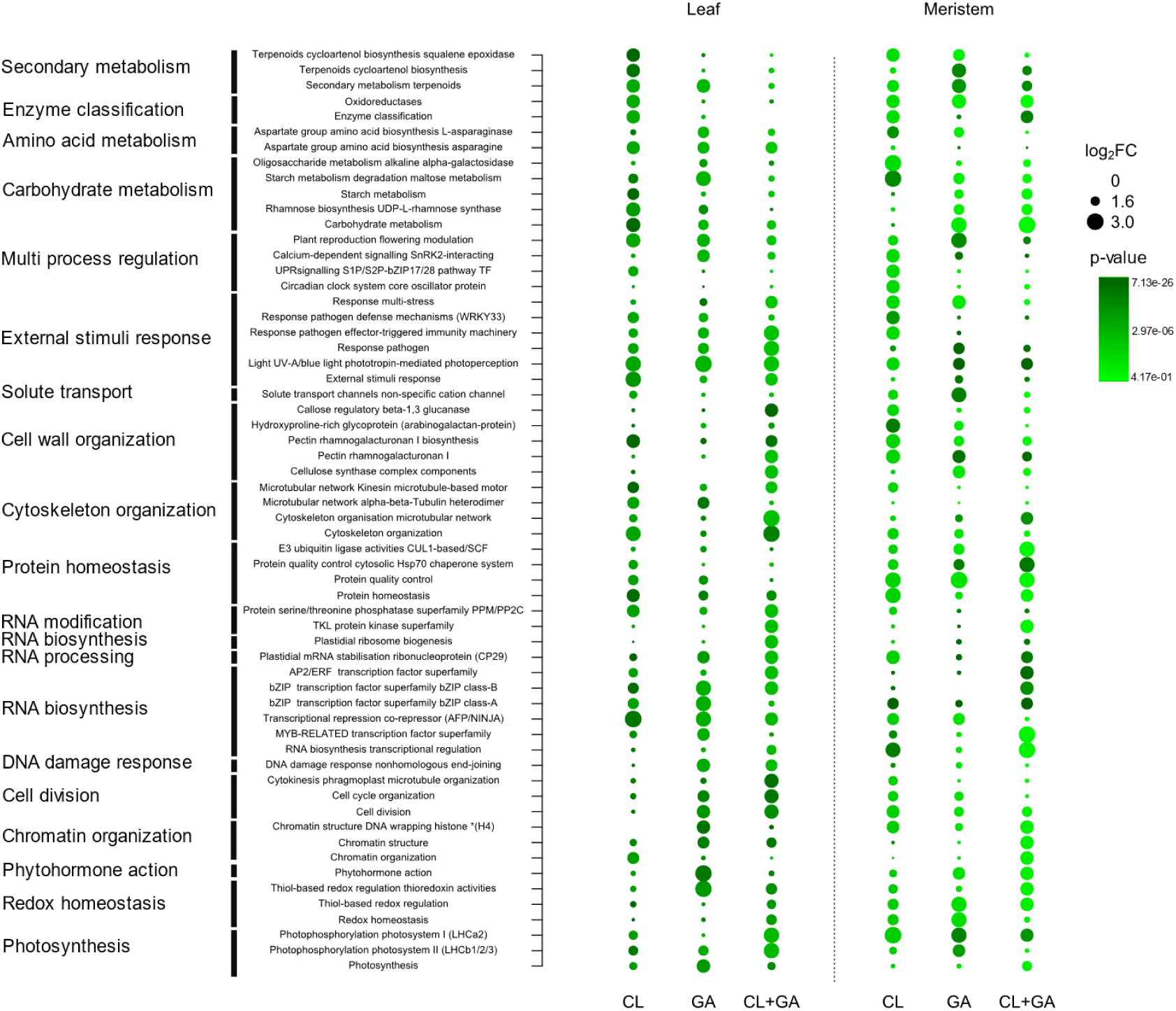
Overrepresentation analysis (ORA) of common genes expressed in the leaf and terminal bud (meristem) tissues (separated by dotted line) of cultivars Clery (CL), Gariguette (GA), or both (CL+GA). Bubble chart shows the log_2_ enrichment factor (log_2_ ERF) for MapMan protein annotation (y-axis) for common genes and the green color gradient represents the adjusted p-value for enriched protein classes.

In the CL terminal bud, the enriched protein classes were similar to those found in leaves, including carbohydrate metabolism, aspartate metabolism, and cell wall organization (hydroxyproline-rich glycoprotein). The DEGs were also enriched for RNA biosynthesis (bZIP superfamily), the circadian clock system, and photosynthesis (Figure 5). In the GA terminal bud, the most enriched protein classes were related to secondary metabolism (terpenoids), responses to external stimuli, cell wall organization (pectin, rhamnogalacturonan), redox homeostasis and photosynthesis (Figure 5). The DEGs shared by both cultivars were enriched for protein classes involved in chromatin organization/structure, cytoskeletal organization (microtubule network), serine/threonine phosphatase superfamily, cell division, cell wall organization (cellulose), RNA biosynthesis (bZIP superfamily) and plant reproduction/flowering (Figure 5).

The geographical location significantly affected the differences observed during floral initiation in both tissues. Protein classes involved in multi-process signaling (phosphoinositides), cytoskeletal organization, and cellular respiration differed most between Germany and France.

### DEGs in early floral initiation highlight known genes involved in the flowering pathway

The common DEGs shared by different genotypes and tissues included those encoding transcription factors and signaling proteins related to plant reproduction and flowering. Some displayed the same tendency regardless of location whereas others showed different behaviors in Germany and France, or in leaves and the terminal bud, or combinations of the location and tissue (Figure 6).

**Figure 6:**
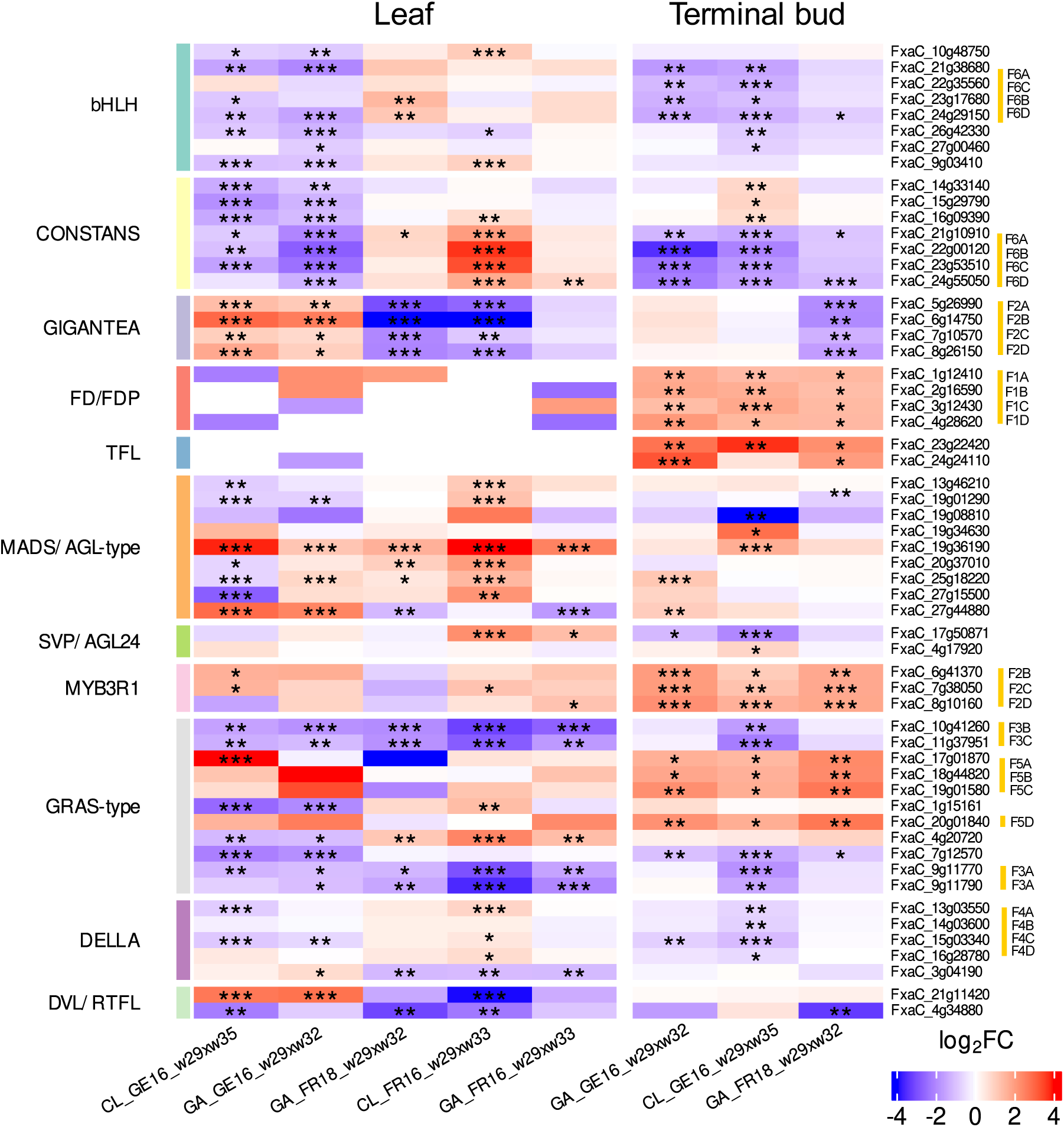
Heat map showing the expression of genes in leaf and terminal bud tissues of strawberry cultivars Clery (CL) and Gariguette (GA) sampled from France (FR) in 2016 and 2018, and from Germany (GE) in 2016, during floral initiation. Differentially expressed genes (named in rows) were chosen based on their protein function (MapMan protein classes) and significance level (***FDR < 0.001, **FDR < 0.01, *FDR < 0.05) for specific date contrasts (shown in columns) during w29 (T0), w32 (T10A), w33 (T10B) and w35 (T50). Gene expression is scaled to range between 5 and –5. Red corresponds to stronger upregulation and blue to downregulation. MapMan protein classification for genes was added on the left and color bars show different protein classes. Homologous genes and their corresponding haplotype in the *F.* × *ananassa* genome are highlighted in yellow on the right.

Seven genes encoding CONSTANS-like (COL) proteins were differentially expressed between T0 and T10 (Figure 7). The downregulation of *COL* genes was observed in all terminal bud samples at T10 compared to T0, with five showing strong, shared downregulation across both cultivars and the other two only significantly downregulated in cultivar GA (Figure 6). In contrast, the differential expression of these *COL* genes in leaves between T0 and T10 was highly dependent on the location. We observed significant downregulation in both cultivars in Germany (CL_GE16_T0xT10L and GA_GE16_T0xT10L) but an increase in France (GA_FR16_T0xT10L and CL&GA_FR18_T0xT10L). The CL_FR16_T0xT10L comparison revealed the significant upregulation of five *COL* genes in leaf tissue.

**Figure 7:**
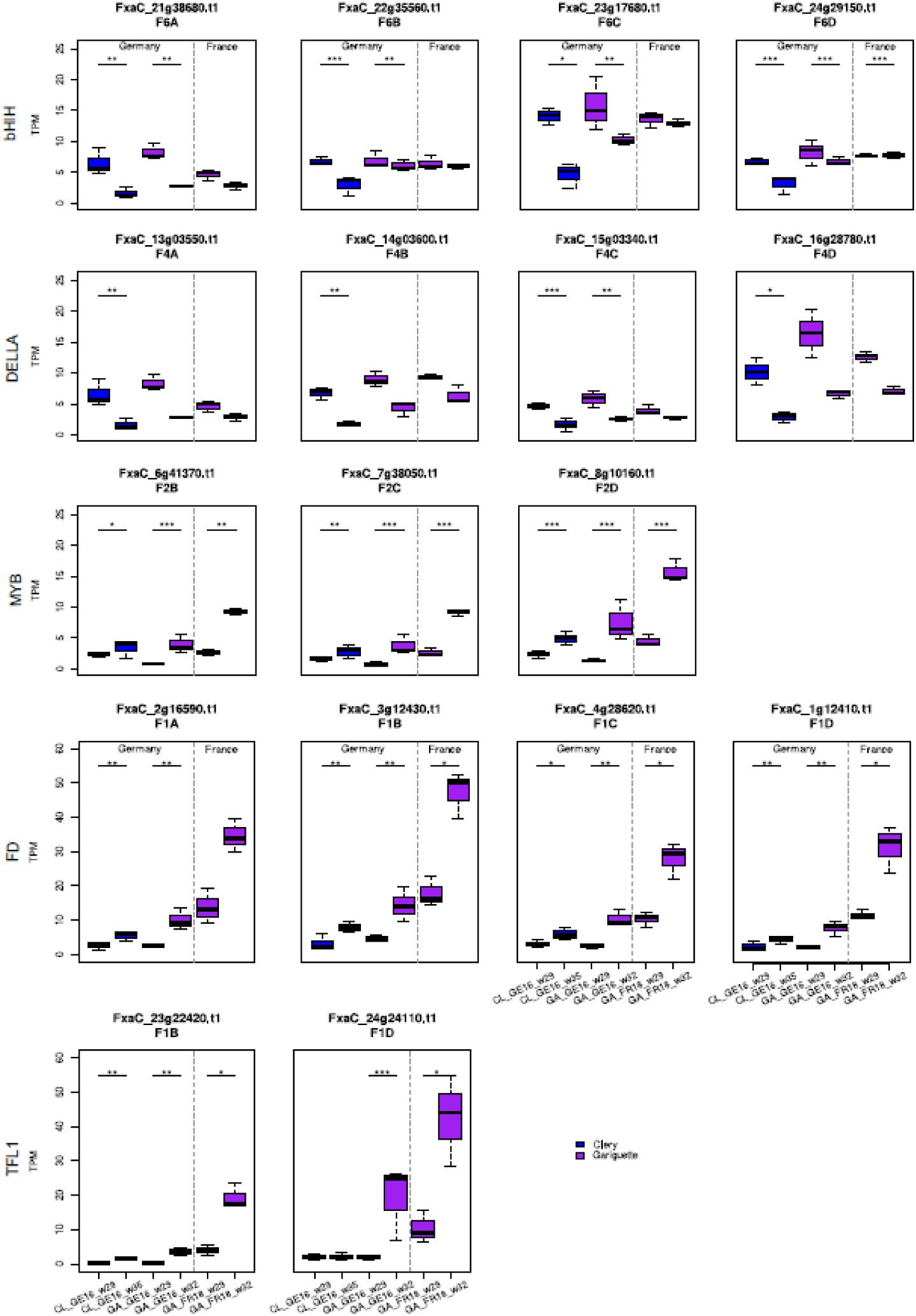
Transcript per million (TPM) values for *bHLH, DELLA, MYB, FD* and *TFL1* candidate genes and their homologous sequences. Comparison of sampling dates during floral initiation in Clery (CL, blue) and Gariguette (GA, purple) terminal buds from plants growing in Germany (GE) and France (FR). Significance levels: ***FDR < 0.001, **FDR < 0.01 and *FDR < 0.05.

Eight genes encoding the bHLH-type transcriptional co-activator FBH were predominantly downregulated between T0 and T10 in the leaf tissue of both cultivars in Germany but tended to be upregulated in France. DEGs encoding the regulatory protein GIGANTEA (GI) were strongly downregulated between T0 and T10 in both tissues of plants grown in France, but were upregulated in leaf samples over the whole floral initiation period in Germany. *FD-like* and *TFL1* genes were minimally expressed in leaves during early floral initiation but were strongly upregulated between T0 and T10 in the terminal buds.

Several MADS/AGL-type genes also showed differential expression. Specifically, the *FxaC_19g36190* gene was significantly upregulated across almost all comparisons in both tissues, except the terminal buds of cultivar GA. Among the DEGs associated with SVP/AGL24, the gene *FxaC_17g50871* was upregulated in the leaves of both cultivars in France but was downregulated in the terminal buds of plants grown in Germany.

DEGs encoding MYB3R1 transcription factors were upregulated in all comparisons of T0 and T10, especially in terminal buds. Transcription factor genes involved in the gibberellin pathway (e.g., encoding DELLA and GRAS-type proteins) were also modulated. Some were significantly downregulated in the leaves of CL plants grown in Germany, and also in the terminal buds of both cultivars, but most such genes in CL_FR16 leaves were upregulated between T0 and T10. Four DEGs encoding GRAS-type proteins were downregulated in all leaf samples, as was one gene in the terminal buds in the case of CL_GE16_ T0xT10, but other DEGs were upregulated in all terminal bud samples. The GA samples in general showed less significant differential expression than CL across locations and years. We identified two DEGs encoding DEVIL/ROT-FOUR-LIKE (DVL/RTFL) proteins with similar expression profiles to *GI* genes. One of these (*FxaC_4g34880*) was significantly downregulated only in the leaves and terminal buds of GA plants grown in France in 2018 (Figure 6).

Given that cultivated strawberry is octoploid (2*n* = 8*x* = 56), up to eight homoeoalleles for each gene located at orthologous positions in the four subgenomes (A, B, C and D) of *F. × ananassa* could potentially be expressed, although only some of the loci may be active (Hardigan et al., 2021). We discuss different situations for different genes below. The expression of homoeoalleles belonging to all four subgenomes was observed in terminal buds for the bHLH transcription factor gene on homoeologous group 6, all of which were significantly downregulated in both cultivars grown in Germany (Figure 7). Similarly, the DELLA transcription factor gene *FanRGA* was expressed in the four subgenomes of homoeologous group 4. They were all significantly downregulated in cultivar CL in Germany, whereas only the alleles of one subgenome (*FxaC_15g03340*) were significantly downregulated in cultivar GA (Figure 7). Finally, the flowering gene *FanFD* was expressed in all four subgenomes whereas *FanTFL1* was only expressed in two subgenomes. These genes were significantly upregulated in most terminal bud samples but no significant differential expression was observed in leaves (Figures 6 and 7; FDR < 0.05).

### Comparative analysis of selected floral initiation genes based on reference data in cultivated strawberry

We compared our DEGs to those reported in a previous study of floral initiation in the cultivated strawberry cv. Benihoppe (Liang *et al*., 2022). This revealed 19 common DEGs in leaf or terminal bud tissues, from which we selected six expressed in the terminal bud and associated with hormone pathways or carbohydrate metabolism for analysis at the subgenomic level (Figure 8).

**Figure 8:**
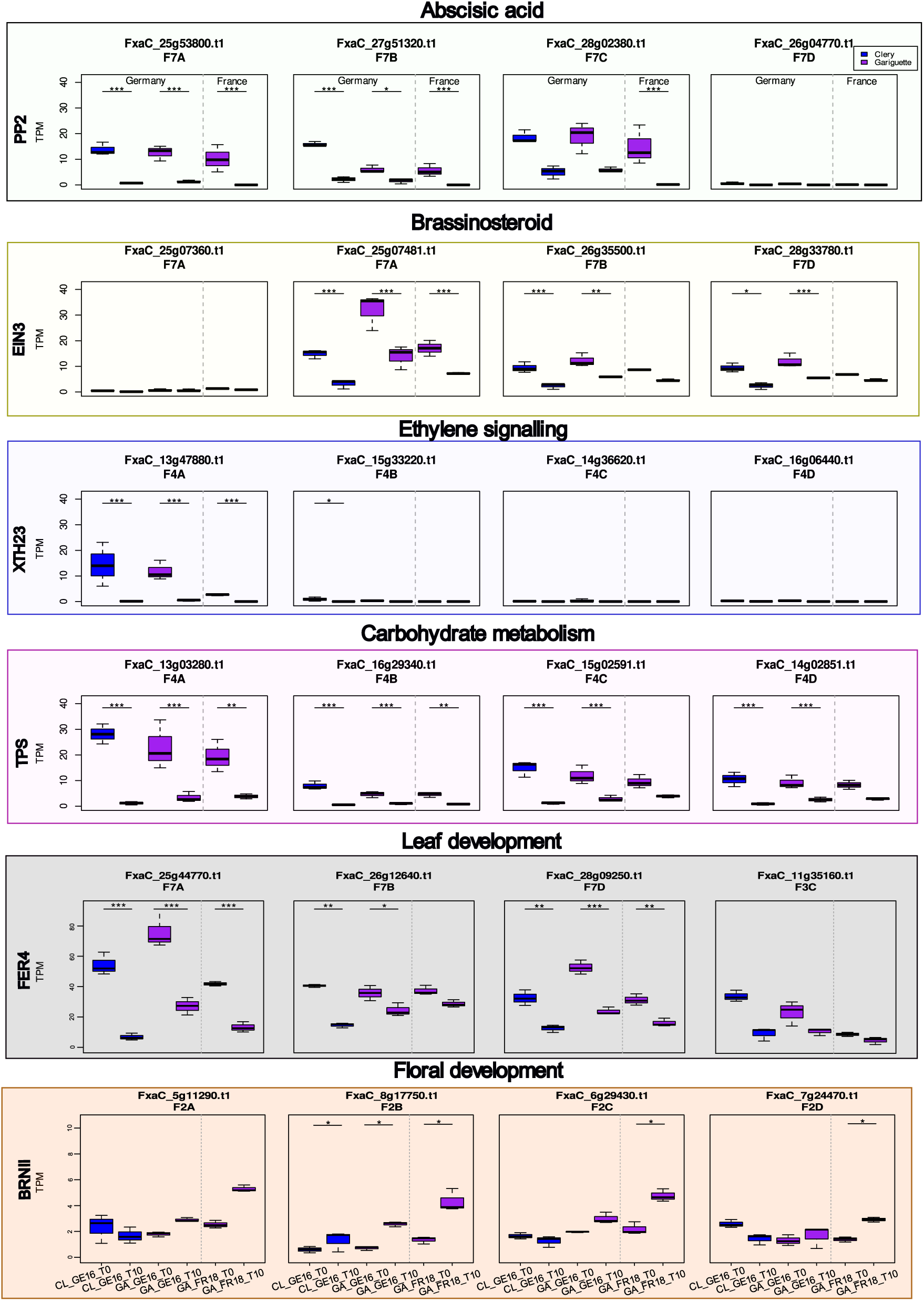
Whisker boxplots illustrating transcript per million (TPM) values for the candidate genes *PP2*, *EIN3*, *XTH23*, *TPS*, *FER4* and *BRNII* involved in the abscisic acid, brassinosteroid and ethylene signaling pathways, and in carbohydrate metabolism. We analyzed the expression of homologous sequences between dates during floral initiation in terminal buds, comparing the Clery (CL, blue) and Gariguette (GA, purple) genotypes growing in Germany (GE) and France (FR). Significance levels: ***FDR < 0.001, **FDR < 0.01 and *FDR < 0.05.

One gene encoding a type 2C protein phosphatase (PP2) involved in the abscisic acid (ABA) pathway was significantly downregulated in three subgenomes across both locations and cultivars, except subgenome 7C (*FxaC_28g02380*) in Germany and subgenome 7D (*FxaC_26g04770*) in the terminal bud (Figure 8). Furthermore, the *EIN3* gene encoding a brassinosteroid ethylene-like transcription factor was downregulated in three subgenomes but strongly expressed in subgenome 7A (*FxaC_25g07481*) in the terminal bud in both Germany and France (Figure 8; FDR < 0.001). The *XTH23* gene involved in ethylene signaling was only significantly downregulated in subgenome 4A for both locations and both cultivars (FDR < 0.001). The expression of *FxaC_15g33220* on subgenome 4B was significantly downregulated for cultivar CL in Germany (Figure 8; FDR < 0.05). In the context of carbohydrate metabolism, a *TPS* gene in homoeology group 4 was expressed at different levels depending on the subgenome, with significant downregulation in subgenomes 4C and 4D only in Germany, but significant downregulation in subgenomes 4A and 4B for both cultivars at both locations (Figure 8).

We analyzed two DEGs associated with in leaf and/or floral development. The *FanFER4* gene, involved in leaf development, was significantly downregulated in both cultivars at both locations (FDR < 0.01) in subgenomes 7A and 7D, but the expression in subgenome 7B was only significant in Germany (FDR < 0.001). Three homeologous alleles of *FER4* were identified on chromosome 7 (FDR < 0.05), and the fourth was located on chromosome 3 (Figure 8). The *BRN1* gene encoding an RNA-binding protein involved in the regulation of flowering time was significantly upregulated in subgenomes 2C and 2D for cultivar GA in France and in subgenome 2B at both locations (Figure 8; FDR < 0.05).

## Discussion

Floral initiation is a highly orchestrated developmental phase influenced by endogenous and exogenous signals (Koorneef et al. 1998; Wellmer and Riechmann, 2010). In cultivated strawberry, a major crop species, the flowering time influences fruit yields, and recent studies have shown that both the genotype and the environment influence floral initiation (Krüger et al., 2022) and flowering time (Prohaska et al., 2024). Floral initiation occurs when temperatures and day length decrease, which reflects interactions between these two environmental factors (Heide et al., 2013). The process therefore varies from year to year at the same location. To accommodate this variation, we based our sampling dates on the percentage of initiated plants rather than a fixed calendar date. By tracking this percentage of initiated plants over time and sampling at 0%, 10% and 50% of initiated plants, we also highlighted the much more rapid floral initiation process in Germany compared to France, probably resulting from the steeper decrease in temperature and day length in Germany than France (Krüger et al., 2022).

The analysis of floral initiation in the model plant Arabidopsis (*Arabidopsis thaliana*) and various crop species has identified several regulatory pathways, including an autonomous pathway as well as those responding to photoperiod, temperature, vernalization, gibberellins, aging and carbohydrate levels (Mouradov et al. 2002; Boss et al., 2004). By studying two cultivars (Gariguette and Clery) cultivated in two different countries (France and Germany) across 3 years, we identified components of these pathways in the regulation of floral initiation in strawberry, as discussed in more detail below.

### Photoperiodic pathway and floral integrators

The photoperiodic pathway of floral initiation involves a network of regulators that respond to day length. In Arabidopsis, the key genes involved in this process are well known. *GI* regulates circadian rhythms and promotes flowering under long-day conditions by stabilizing the CO protein (Michaels et al., 2003; Brandoli et al., 2020). CO plays a central role in sensing day length, accumulating during long days to directly activate the expression of the florigen gene *FT* (Kardailsky et al., 1999; Kobayashi et al., 1999; Abe et al., 2005; Wigge et al., 2005; Corbesier et al., 2007). In the SAM, FT competes with the floral repressor TFL1 to bind FD (Zhu et al. 2020). SOC1 is an integrator of the photoperiod, gibberellin and temperature pathways to promote flowering at the shoot apex (Li et al., 2008; Jung et al., 2012).

We found that the strawberry orthologs of these genes (*FanCO*, *FanSOC1*, *FanFD*, *FanTFL1* and *FanGI*) were differentially expressed during early floral initiation. The involvement of *FanCO*, *FanSOC1* and *FanTFL1* in the seasonal control of floral initiation has already been demonstrated (Iwata et al. 2012; Koskela et al. 2012; Mouhu et al. 2013; Koskela et al. 2016; Korukura et al. 2017; Munoz-Avila et al. 2022). In addition, we and others (Liang et al. 2022) showed that *FanBRN1* is also differentially expressed during floral initiation in strawberry. The Arabidopsis ortholog *AtBRN1* acts as a repressor of SOC1 activity (Kim et al., 2013).

### Phytohormone signaling

We identified multiple DEGs associated with the gibberellic acid, brassinosteroid, ABA, and jasmonic acid pathways. The gibberellic acid pathway promotes the transition from vegetative growth to flowering (Yu et al., 2004; Mutasa-Göttgens & Hedden 2009; Achard & Genschik, 2009). We found that *FanRGA* was differentially expressed in the terminal buds of both cultivars. In diploid strawberry, the ortholog *FveRGA* encodes a DELLA protein that suppresses stolon development (Li et al. 2018; Caruana et al. 2018). DELLA proteins are negative regulators of gibberellin signaling that act immediately downstream of the gibberellic acid receptor (Eckardt 2007).

Brassinosteroids promote flowering in Arabidopsis, (Liz and He, 2020). We found that *FanXTH2,* a component of the brassinosteroid pathway encoding the enzyme xyloglucan endotransglucosylase/hydrolase, was differentially expressed between cultivars and locations, as previously shown in strawberry terminal bud tissue (Liang et al., 2022) and loquat bud tissue (Xia et al., 2020).

We observed the downregulation of *FanPP2* during floral initiation in both cultivars and locations, as also reported in a previous study (Liang et al., 2022). The Arabidopsis ortholog encodes a negative regulator of ABA signaling, suggesting that it influences the timing of floral initiation by repressing ABA-responsive genes. We also observed the downregulation of *FanEIN3*, encoding an ethylene-responsive transcription factor, consistently with previous findings (Liang et al., 2022). The Arabidopsis ortholog delays flowering by activating *ERF1* and the APETALA2 (AP2)/ERF1 protein family (Guo and Ecker, 2004).

### Transcription factors

Multiple transcription factor families, including bHLH, MADS-box and MYB, were differentially expressed during floral initiation, and are known to be involved in the transition from vegetative growth to reproductive development (Bemer & Angenent 2009; Woodger et al., 2003; Liang et al., 2022). For example, we found two differentially expressed members of the DVL/RTFL family, which are known to be involved in organogenesis (Guo et al., 2015). One was upregulated in both cultivars and locations, but the other was downregulated in German samples and upregulated in French samples of the cultivar CL. Similarly, Arabidopsis DVL/RTFL proteins influence traits such as leaf shape and plant architecture. The overexpression of *DEVIL1* (*DVL1*) and *ROTUNDIFOLIA4* (*ROT4/DVL16*) in Arabidopsis produced a pleiotropic phenotype with short stature, rounder rosette leaves, and clustered inflorescences (Wen et al., 2004; Valdivia et al., 2012).

Plant peptides also play important roles in development, regulating terminal bud organization, root growth, and leaf shape (Matsubayashi 2014; Matsubayashi and Sakagami 2006; Guo et al., 2015). We observed the tissue-specific expression of GRAS-type proteins, which are putative transcriptional regulators, consistent with their documented roles. Most GRAS-type genes were dominantly expressed in roots with a subset also expressed in shoots and leaves (Heckmann et al., 2006; Liang et al., 2022). The findings emphasize the interplay between transcriptional regulation and tissue-specific signaling during floral initiation.

### Carbohydrate and energy metabolism

DEGs related to carbohydrate (particularly starch) metabolism were identified in both cultivars, as well as certain secondary metabolic pathways, such as the terpenoid pathway (Figure 5). Trehalose-6-phosphate (T6P) is a sugar derivative that is proposed to influence flowering both in the terminal bud and the SAM. Accordingly, the *TPS1* gene encoding trehalose-6-phosphate synthase contributes to the genetic framework that controls flowering time (Wahl et al., 2013; Rojas et al., 2023). We showed that *FanTPS1* was downregulated in terminal buds in both cultivars and locations, as previously reported (Liang et al., 2022).

The carbohydrate pathway not only controls floral initiation but also contributes more widely to plant development (Zhang et al., 2013; Qiao et al., 2021). Carbohydrate metabolism provides the energy and building blocks needed for flower development so the interplay between carbohydrate metabolism, phytohormonal signaling and environmental cues ensures that the transition from vegetative growth to flowering is timed to maximize reproductive success (Corbesier et al., 1998; Sawicki et al., 2015). In the terminal bud of cultivar CL, we observed the overrepresentation of DEGs linked to starch metabolism, which serves as an energy reserve. Mobilized starch, converted into sucrose in leaves and stems, provides an early signal for floral induction (Bernier et al., 1993). Key differences between the cultivars included higher asparagine and rhamnogalacturonan I (RG-I) production in GA. Asparagine provides nitrogen for signaling proteins that coordinate floral induction with internal and external pathways (Khurana et al., 1988).

The *FERONIA 4 (FER4*) gene encodes a product that contributes to carbohydrate metabolism (particularly glycolysis) by interacting with the cytosolic enzyme glyceraldehyde-3-phosphate dehydrogenase (GAPDH). *FER* deficiency has been shown to reduce GAPDH activity, causing the of accumulation starch (Yang et al., 2015). We found that *FER4* was downregulated in both cultivars, and that protein classes related to starch metabolism were enriched. *FER4* was also downregulated in the cultivated strawberry cultivar Benihoppe (Liang et al., 2022), where it was linked to leaf development.

### Cytoskeleton, cell division and cell wall dynamics

In the terminal buds of cultivar GA, we identified DEGs associated with cytoskeletal organization, specifically the microtubule network required for cell division and expansion, thereby influencing tissue patterning during floral initiation (Chandler 2012; Denay et al., 2017). Interestingly, DEGs related to cell division were overrepresented in the terminal buds of cultivar CL, whereas DEGs related tRNA biosynthesis, bZIP transcription factors and RG-I biosynthesis, the latter needed for cell wall integrity (Yapo, 2011), were overrepresented in the terminal buds of cultivar GA. The role of RG-I in floral induction remains unclear. Disparities in the representation of protein classes between cultivars may reflect the field experiment setup and environmental fluctuations, masking inherent differences between the cultivars.

### Regulation of subgenomic expression

In the octoploid cultivated strawberry, the plasticity of traits such as flowering (Prohaska et al., 2024) can confer polyploid advantage in heterogeneous environments (Wei et al., 2019). In this species, each gene may be represented by up to eight homoeoalleles located on the four homoeologous subgenomes (Rousseau-Gueutin et al., 2019). However, polyploidization is followed by a process of diploidization, whereby gene redundancy is reduced by processes such as gene silencing, sequence elimination and rearrangement (Chen 2007; Doyle et al. 2008). In strawberry, the analysis of gene redundancy revealed that only 46% of genes retain alleles in all four subgenomes, whereas 7%, 23% and 14% retain alleles on one, two and three homoeologous chromosomes, respectively (Jin et al., 2023). For most of the *F. × ananassa* genes we identified, such as those encoding FBH, CO, GI, GRAS-type, DELLA, FD/FDP, PP2, XTH23, TPS and BRN1 proteins, we found sequences in all four subgenomes, whereas *MYB3R1*, *EIN3* and *FER4* sequences were present in three subgenomes and were similarly expressed. The expression of these genes is therefore likely to be finely regulated, with mutations in the promoter and/or 5′ untranslated region.

## Conclusion

Our study sheds light on the molecular basis of early floral initiation in *F. × ananassa* under natural conditions. We identified key gene expression profiles and differences between cultivars CL and GA across two environments (Germany and France). We highlighted the roles of genes such as *XTH23, TPS and FER4*, and those encoding transcription factors such as FBH, CO and GI, in the regulation of this developmental transition, which also involves the reprogramming of carbohydrate metabolism, phytohormone signaling, and photoperiod-dependent regulation. These findings offer valuable insights for the optimization of strawberry cultivation and breeding strategies, but more research is needed to determine the precise functions of the key genes involved in floral initiation and the influence of their differential expression.

## Materials and methods

### Floral initiation

The timing of floral initiation was determined by non-invasive architectural analysis of the *Fragaria × ananassa* cultivars Gariguette (GA) and Clery (CL), discriminating between the vegetative and floral stages of the SAM in the terminal bud. Plant architecture assessments were conducted over time throughout the summer and autumn at two locations: Bordeaux in France and Geisenheim in Germany. The first samples were collected in mid-July (T0, w29), coinciding with the presence of daughter plants featuring only small root primordia on stolons. Subsequent sampling took place in August (w32), 3 weeks after the transplantation of rooted plants to the field or nursery, and at 10-day intervals until early October (w40), making eight sampling dates in total.

Plant architecture was assessed as previously described (Krüger et al., 2022). Briefly, we described the daughter plants (including the number of developed leaves and the stage of the terminal bud) using a stereomicroscope with 40–60× magnification. A single terminal bud from the main crown of each plant was dissected for analysis (Figure 2A). We dissected 9–12 plants representing each cultivar and environment at each sampling date. The vegetative or floral status of the terminal buds was assessed as previously described (Krüger et al., 2022) as adapted from earlier methods (Jahn & Dana 1970; Taylor et al., 1997).

### RNA-Seq sampling

RNA-Seq sampling time points were determined by the percentage of initiated SAM. Based on the assumption that samples at 0% (time point T0) and 5–20% (time point T10) of initiated plants would capture early floral initiation steps, samples of leaves and terminal buds were collected accordingly at three or four dates depending on the location and year. For each date, cultivar and organ, we collected three replicate samples of nine leaf discs or nine terminal buds from nice separate plants in one 1.5-ml Eppendorf tube (Figure 2B). RNA was extracted as previously described (Gaston et al., 2021) and 3 µg per sample was sent to Sistemas Genomicos (Spain) for sequencing on the Illumina HiSeq 2500 platform.

### RNA-Seq data processing

Adaptors were removed from the Illumina RNA-Seq dataset using Trimmomatic v0.38 (Bolger et al., 2014) followed by alignment to the Fragaria_ananassa_v1.0.a2 cv. Camarosa reference genome (GDR database https://www.rosaceae.org; Liu et al. 2021) using the pseudo-aligner Salmon v1.10.1 for read count quantification (Patro et al., 2017). DEGs were identified using edgeR v3.40.1 (Robinson et al., 2010). Contrasts were calculated between the different sampling points (T0, T10A, T10B and T50) and cultivars separately for both tissues. The transcripts per million (TPM) were evaluated by PCA, and ORA was applied to the PC1 and PC2 axes. This provided insights into the leading MapMan (Schwacke et al., 2019) protein classes contributing to the separation along PC1 and PC2 in the positive and negative directions. The outcomes were visually represented as bar charts illustrating the enrichment factor along these axes. The top five genes in both directions along PC1 and PC2 were also depicted on the PCA plot. Venn diagrams were generated to identify shared DEGs in different comparisons using VennDiagram v1.7.3 (Chen & Boutros, 2011) in R. A threshold of FDR < 0.05 was used to filter significant DEGs. Heat map clustering was applied to discern gene relationships within each tissue.

### Gene correspondence and functional analysis

To link our work to the published literature (Li et al., 2019; Liang et al., 2022), we matched gene names between the current Camarosa *F. × ananassa* annotation (v2) and the diploid genome of *F. vesca* or the Camarosa v1 annotation used by Liang et al. (2022). The genes were checked bidirectionally against the diploid transcriptome of *F. vesca* and *F. × ananassa* cv. Camarosa using Blast+ v2.15.0 (Camacho *et al*., 2009). We focused on the DEGs in our study and applied ORA to the overlapping genes to identify MapMan protein classes that potentially influence floral initiation. Statistical analysis was carried out using R Studio base version 2023.06.0 (RStudio Team, 2023).

## Declarations

### Ethics approval and consent to participate

Not applicable

### Consent for publication

Not applicable

### Availability of data and materials

Raw data will be available on EMBL-EBI (PRJEB83679).

## Acknowledgement

This study was funded by the European Union’s Horizon 2020 program under grant agreement number 679303. We thank Dr. Richard M Twyman for manuscript proofreading.

## Contributions

FMRZ analyzed the data and wrote the manuscript, supported and supervised by AG, BD and BU. AG, BD, MD, KG, SO, EK and BB generated and provided RNA-Seq data. All authors read and approved the final manuscript.

## References

Abe M, Kosaka S, Shibuta M, Nagata K, Uemura T, Nakano A, Kaya H (2019). Transient activity of the florigen complex during the floral transition in Arabidopsis thaliana. Plant Development 146(7):dev171504.

Achard P, Baghour M, CHapple A, Hedden P, Van der Straeten D, Genschik P, Moritz T, Harberd NP (2007). The plant stress hormone ethylene controls floral transition via DELLA-dependent regulation of floral meristem-identity genes. PNAS 104(15): 6484–6489.

Achard P & Genschik P (2009). Releasing the brakes of plant growth: how GAs shutdown DELLA proteins. Journal of Experimental Botany 60(4). 1085–1092.

Bao S, Hua C, Shen L, Yu H (2019). New insights into gibberellin signaling in regulating flowering in Arabidopsis. Journal of Integrative Plant Biology 62(1): 118–131.

Boss PK, Bastow RM, Mylne JS, Dean C (2004). Multiple pathways in the decision to flower: enabling, promoting, and resetting. Plant Cell. 16.

Battey NH, Le Mière P, Tehranifar A, Cekic C, Taylor S, Shrives KJ, Hadley P, Greenland AJ, Darby J, Wilkinson MJ (1998). Genetic and environmental control of flowering in strawberry. Genetic and environmental manipulation of horticultural crops: CAB international 111–131.

Bemer M & Angenent GC (2009). Floral organ initiation and development. Plant Developmental Biology-Biotechnological Perspectives 173–194.

Benlloch R, Berbel A, Serrano-Mislata A, Madueño F (2007). Floral initiation and inflorescence architecture: a comparative view. Annals of Botany 100(3): 659–676.

Bernier G, Havelange A, Houssa C, Petitjean A, Lejeune P (1993). Physiological signals that induce flowering. Plant Cell 5(10). 1147–1155.

Bolger AM, Lohse M, Usadel B (2014). Trimmomatic: a flexible trimmer for Illumina sequence data. Bioinformatics 30(15): 2114–2120.

Brandoli C, Petri C, Egea-Cortines M, Weiss J (2020). Gigantea: Uncovering new functions in flower development. Genes 11(10): 1142.

Caruana JC, Sittman JW, Wang W, Liu Z (2018). *Suppressor of Runnerless* Encodes a DELLA protein that controls runner formation for asexual reproduction in strawberry. Molecular Plant 11: 230–233.

Camacho C, Coulouris G, Avagyan V, Ma N, Papadopoulos J, Bealer K, Madden TL (2009). BLAST+: architecture and applications. BMC Bioinformatics 10: 421.

Cheng H, Qin L, Lee S, Fu X, Richards DE, Cao D, Luo D, Harberd NP, Peng J (2004). Gibberellin regulates Arabidopsis floral development cis suppression of DELLA protein function. 131(5): 1055–1064.

Chandler JW (2012). Floral meristem initiation and emergence in plants. Cellular and Molecular Life Sciences 69: 3807–3818.

Chen ZJ (2007) Genetic and epigenetic mechanisms for gene expression and phenotypic variation in plant polyploids. Annual Review of Plant Biology 58:377–406.

Chen H & Boutros PC (2011). VennDiagram: a package for generation of highly customizable Venn and Euler diagrams in R. BMC Bioinformatics 12(35).

Corbesier L, Lejeune P, Bernier G (1998). The role of carbohydrates in the induction of flowering in *Arabidopsis thaliana*: comparison between the wild type and a starchless mutant. Planta 206: 131–137.

Costes E, Crespel L, Denoyes B, Morel P, Demene MN, Lauri PE, Wenden B (2014). Bud structure, position and fate generate various branching patterns along shoots of closely related Rosaceae species: a review. Frontiers in Plant Science 5.

Darrow GM (1996). The Strawberry. CAB Direct. ref.bibl.230; pp. 447 pp.

Davis SJ (2009). Integrating hormones into floral-transition pathway of *Arabidopsis thaliana*. Plant, Cell & Environment 32(9): 1201–1210.

Denay G, Chahtane H, Tichtinsky G, Parcy F (2017). A flower is born: an update on Arabidopsis floral meristem formation. Current Opinion in Plant Biology 35: 15–22.

Durner EF, Barden JA, Himelrick DG, Poling EB (1984). Photoperiod and temperature effects on flower and runner development in day-neutral, junebearing, everbearing strawberries. Journal of the American Society for Horticultural Science 109(3): 396–400.

Doyle JJ, Flagel LE, Paterson AH, Rapp RA, Soltis DE, Soltis PS, Wendel JF (2008). Evolutionary Genetics of Genome Merger and Doubling in Plants. Annual Review in Genetics 42:443–461.

Eckhardt NA (2007). GA signaling: direct targets of DELLA proteins. The Plant Cell 19(10): 2970.

Fu Y (2010). The actin cytoskeleton and signaling network during pollen tube tip growth. Journal of Integrative Plant Biology 52(2): 131–137.

Gaston A, Perotte J, Lerceteau-Köhler E, Rousseau-Guetin M, Petit A, Hernould M, Rothan C, Denoyes B (2013). PFRU, a single dominant locus regulates the balance between sexual and asexual plant reproduction in cultivated strawberry. Journal of Experimental Botany 64(7): 1837–1848.

Gaston A, Potier A, Alonso, Sabbadini S, Delmas F, Tenreira T, Cochetel N, Labadie M, Prévost P, Folta KM, Mezzeti B, Hernould M, Rothan C, Denoyes B (2021). The *FveFT2* florigen/*FveTFL1* antiflorigen balance is critical for the control of seasonal flowering in strawberry while *FveFT3* modulates axillary meristem fate and yield. New Phytologist 232(1): 372–387.

Gossens J, Mertens J, Gossens A (2017). Role and functioning of bHLH transcription factors in jasmonate signalling. Journal of Experimental Botany 68(6): 1333–1347.

Guo HW and Ecker JR (2004). The ethylene signaling pathway: New insights. Current Opinion Plant Biology 7:40–49.

Guo P, Yoshimura A, Ishikawa N, Yamaguchi T, Guo Y, Tsukaya H (2015). Journal of Plant Research 128(3): 497–510.

Hancock JF, Luby JJ, Dale A, Callow PW, Serçe S, El-Shiek A (2002). Utilizing wild *Fragaria virginiana* in strawberry cultivar development: Inheritance of photoperiod sensitivity, fruit size, gender, female fertility and disease resistance. Euphytica 126: 177–184.

Heckmann AB, Lombardo F, Miwa H, Perry JA, Bunnewell S, Parniske M, Wang TL, Downie JA (2006). Lotus japanicus nodulation requires two GRAS domain regulators, one which is functionally conserved in a non-legume. Plant Physiology 142(4): 1739–1750.

Heide OM, Stavang JA, Sønsteby A (2013). Physiology and genetics of flowering in cultivated and wild strawberries-a review. The Journal of Horticultural Science and Biotechnology 88(1).

Hytonen T and Kurokura (2020). Control of flowering and runnering in Strawberry. The Horticulture Journal 89(2):96–107.

Hung FY, Lai YC, Wang J, Feng YR, Shih YH, Chen JH, Sun HC, Yang S, Li C, Wu K (2021). The Arabidopsis histone demethylase JMJ28 regulates CONSTANS by interacting with FBH transcription factors. The Plant Cell 33(4): 1196–1211.

Ito S, Song YH, Joseph-Day AR, Miller RJ, Breton G, Olmsted RG, Imaizumi T (2012). FLOWERING BHLH transcriptional activators control expression of the photoperiodic flowering regulator CONSTANS in Arabidopsis. PNAS 109(9): 3582–3587.

Iwata H, Gaston A, Remay A, Thouroude T, Jeauffre J, Kawamura K, Oyant LHS, Araki T, Denoyes B, Foucher F (2011). The TFL1 homologue KSN is a regulator of continuous flowering in rose and strawberry. The Plant Journal 69(1):116–125.

Jin X, Du H, ZhuC, Wan H, Liu F, Ruan J, Mower JP, Zhu A (2023). Haplotype-resolved genomes of wild octoploid progenitors illuminate genomic diversifications from wild relatives to cultivated strawberry. Nature Plants 9: 1252–1266.

Kawamoto N, Sasabe M, Endo M, Machida Y, Araky Y (2015). Calcium-dependent protein kinases responsible for the phosphorylation of a bZIP transcription factor FD crucial for the florign complex formation. Nature Scientific reports 5: 8341.

Khurana JP, Tamot BK, Maheshwari SC (1988). Floral induction in a photoperiodically insensitive duckweed, *Lemna paucicostata* LP6: Role of glutamate, aspartate, and other amino acids and amides. Plant Physiology 86(3): 904–907.

Kim HS, Abbasi N, Choi SB (2013). Bruno-like proteins mod late lowering time via ′ U R-dependent decay of SOC1 mRNA. New Phytologist 198(3): 747–756.

Koembuoy K, Hasegawa S, Otagaki S, Takahashi H, Nagano S, Isobe S, Shiratake K, Matsumoto S (2020). RNA-Seq analysis pf meristem cells indentifies the FaFT3 gene as a common floral inducer in japaneses cultivated strawberry. The Horticulture Journal 89(2): 138–146.

Korneef M, Alonso-Blanco Cm, Peeters JM, Soppe W (1998). Genetic control of flowering time in Arabidopsis. Annual Review of Plant Biology 49: 345–370.

Koskela EA, Mouha K, Albani MC, Kuokura T, Rantanen M, Sargent DJ, Battey NH, Coupland G, Elomaa P, Hytönen T (2012). Mutation in TERMINAL FLOWER 1 reverses the photoperiodic requirement for flowering in the wild strawberry Fragaria vesca. Plant Physiology 159(3): 1043–1054.

Koskela EA, Sonsteby A, Flachowsky H, Heide Om, Hanke MV, Elomaa P, Hytönen T (2016). TERMINAL FLOWER 1 is a breeding target for a novel everbearing trait and tailored flowering responses in cultivated strawberry (Fragaria x ananassa Duch.). Plant Biotechnology Journal 14(9): 1852–1861.

Krüger E, Woznicki TL, Heide OM, Kusnierek K, Rivero R, Masny A, Sowik I, Brauksiepe B, Eimert K, Mott D, Savini G, Demene M, Guy K, Petit A, Denoyes B, Sonsteby A (2022). Flowering Phenology of Six Seasonal-Flowering Strawberry Cultivars in a Coordinated European Study. Horticulturae 8(10): 933.

Kurokura T, Samad S, Koskela E, Mouhu K, Hytönen T (2017). *Fragaria vesca* CONSTANS controls photoperiodic flowering and vegetative development. Journal of Experimental Botany 68(17): 4839–4850.

Langridge J (1957). Effect of day-length and gibberellic acid on the flowering of Arabidopsis thaliana. Nature 180: 36–37.

Liang J, Zheng J, Wu Z, Wang H (2022). Time-course transcriptomic profiling of floral induction in cultivated strawberry. International Journal of Molecular Sciences 23: 6126.

Liu T, Li M, Liu Z, Ai X, Li Y (2021). Reannotation of the cultivated strawberry genome and establishment of a strawberry genome database. Horticulture Research 8:41.

Matsubayashi Y, Sakagami Y (2006). Peptide hormones in plants. Annual Reviews in Plant Biology 57: 649–674.

Matsubayashi Y (2014). Posttranslationally modified small-peptides signals in plants. Annual Reviews in Plant Biology 65: 385–413.

Mezetti B, Hernould M, Rothan C, Denoyes B (2021). The FveFT2 florigen/FveTFL1 antiflorigen balance is critical for the control of seasonal flowering in strawberry while FveFt3 modulates axillary meristem fate and yield. New Phytologist 232(1): 372–387.

Michaels SD, Ditta G, Gustafson_Brown C, Pelaz S, Yanofsky M, Amasino RM (2003). AGL24 acts as promoter of flowering in Arabidopsis and is positively regulated by vernalization. The Plant Journal 33(5): 867–874.

Moon J, Suh SS, Lee H, Choi KR, Hong CB, Paek NC, Kim SG, Lee I (2003). The SOC1 MADS-box gene integrates vernalization and gibberellin signals for flowering in *Arabidopsis*. The Plant Journal 35(5): 613–623.

Mouradov A, Cremer F, Coupland G (2002). Control of flowering time: Interacting pathways as a basis for diversity. Plant Cell 14: 111–130.

Mutasa-Göttgens E & Hedden P (2009). Gibberellin as a factor in floral regulatory networks. Journal Experimental Botany 60(7): 1979–1989.

Oritz-Marchena MI, Albi T, Lucas-Reina E, Said FE, Romero-Campero FJ, Cano B, Ruiz MT, Romero JM, Valverde F (2014). Photoperiodic control of carbon distribution during the floral transition in Arabidopsis. The Plant Cell 26(2): 565–584.

Patro R, Guggal G, Love MI, Irizarry RA, Kingsford C (2017). Salmon: fast and bias-aware quantification of transcript expression using dual-phase inference. Nature Methods 14(4): 417–419.

Perilleux C, Bouche F, Randoux M, Orman-Ligeza B (2019). Turning meristems into fortresses. Trends in Plant Science 24: 431–442.

Prohaska A, Petit A, Lesemann S, Rey-Serra P, Mazzoni L, Masny A, Sánchez-Sevilla JF, Potier A, Gaston A, Klamkowski K, Rothan C, Mezzetti B, Amaya I, Olbricht K, Denoyes B (2024). Strawberry phenotypic plasticity in flowering time is driven by the interaction between genetic loci and temperature. Journal Experimental Botany.75(18):5923–5939.

Quiao Z, Hu H, Shi S, Yuan X, Yan B, Chen L (2021). An update on the function, biosynthesis and regulation of floral volatile terpenoids. Horticulturae 7(11): 451.

Rantanen M, Kurokura T, Mouhu K, Pinho P, Tetri E, Halonen L, Palonen P, Elomaa P, Hytönen T (2014). Light quality regulates flowering in FvFT1/FvTFL1 dependent manner in the woodland strawberry Fragaria vesca. Frontiers in Plant Science 5.

Robinson MD, McCarthy DJ, Smyth GK (2010). edgeR: a Bioconductor package for differential expression analysis of digital gene expression data. Bioinformatics 26(1): 139–140.

Rojas BE, Tonetti T, Figueroa CM (2023). Trehalose 6-phosphate metabolism in C4 species. Current Opinion in Plant Biology 72.

Rousseau-Gueutin M, Gaston A, Aïnouche A, Aïnouche ML, Olbricht K, Staudt G, Richard L, Denoyes-Rothan B (2009). Tracking the evolutionary history of polyploidy in Fragaria L. (strawberry): new insights from phylogenetic analyses of low-copy nuclear genes. Molecular Phylogenetics Evolution 51(3):515–30.

Savini G, Neri D, Zucconi F, Sugiyama N (2005). Strawberry Growth and Flowering, International Journal of Fruit Science 5(1): 29–50.

Sawicki M, Jacquens L, Baillieul F, Clément C, Vaillant-Gaveau N, Jacquard C (2015). Distinct regulation in inflorescence carbohydrate metabolism according to grapevine cultivars during floral development. Physiologia Plantarum 154(3): 447–467.

Schwacke R, Ponce-Soto GY, Krause K, Bolger AM, Arsova B, Hallab A, Gruden K, Stitt M, Bolger ME, Usadel B (2019). MapMan4: A refined protein classification and annotation framework applicable to multi-omics data analysis. Molecular Plant 12(6): 879–892.

Stewart PJ & Folta KM (2010). A review of photoperiodic flowering research in strawberry (*Fragaria* spp.). Critical Reviews in Plant Sciences 29(1): 1–13.

van Schie CCN, Haring MA, Schuurink RC (2006). Regulation of terpenoid and benzoid production in flowers. Current Opinion in Plant Biology 9(2): 203–208.

Sønsteby A & Heide OM (2008). Temperature responses, flowering and fruit yield of the June-bearing strawberry cultivars Florence, Frida and Korona. Scientia Horticulturae 119(1): 49–54.

Tenreira T, Lange MJP, Lange T, Bres C, Labadie M, Monfort A, Hernould M, Rothan C, Denoyes B (2017). A specific gibberellin 20-oxidase dictates the flowering-runnering decision in diploid strawberry. The Plant Cell 29(9): 2168–2182.

Valdivia ER, Chevalier D, Sampedro J, Taylor I, Niederhuth CE, Walker JC (2012). DVL genes play a role in the coordination of socket cell recruitment and differentiation. Journal of Experimental Botany 63(3): 1405–1412.

Verheul MJ, Sønsteby A, Grimstad SO (2006). Interactions of photoperiod, temperature, duration of short-day treatment and plant age on day treatment and plant age on flowering of *Fragaria* x *ananassa* Duch. cv. Korona. Scientia Horticulturae 107(2): 164–170.

Verheul MJ, Sønsteby A, Grimstad SO (2007). Influences of day and night temperatures on flowering of *Fragaria* x *ananassa* Duch., cvs. Korona and Elsanta, at different photoperiods. Scientia Horticulturae 112(2): 200–206.

Vince-Prue D, Guttridge CG (1973). Floral initiation in strawberry: Spectral evidence for the regulation of flowering by long-day inhibition. Planta 110: 165–172.

Wahl V, Punnu J, Schlereth A, Arrivault S, Langenecker T, Franke A, Feil R, Lunn JE, Stitt M, Schmid M (2013). Regulation of Flowering by Trehalose-6-Phosphate Signaling in Arabidopsis thaliana. Science 339(6120): 704–707.

Wang W, Sijacic P, Xu P, Lian H, Liu Z (2018). *Arabidopsis* TSO1 and MYB3R1 from regulatory module to coordinate cell proliferation with differentiation in shoot and root. PNAS 115(13): E3045–E3054.

Weberling F (1989). Morphology of flowers and inflorescences. 1st Edition Cambridge, UK Cambridge University Press.

Wei N, Cronn R, Liston A, Ashman TL (2019). Functional trait divergence and trait plasticity confer polyploid advantage in heterogeneous environments. New Phytologist 221(4):2286–2297

Wen J, Lease KA, Walker JC (2004). DVL, a novel class of small polypeptides: overexpression alters Arabidopsis development. The Plant Journal. 37(5): 668–677.

Wellmer F, Riechmann JL (2010). Gene networks controlling the initiation of flower development. Review Trends in Genetics 26(12): 519–527.

Wickland DP, Hanzawa Y (2015). The FLOWERING LOCUS TERMINAL FLOWER 1 Gene Family: Functional evolution and molecular mechanisms. Molecular Plant 8; 983–997.

Woodger FJ, Millar A, Murray F, Jacobson JV, Gubler F (2003). The role of GAMYB transcription factors in GA-regulated gene expression. Journal of Plant Growth Regulation. 22: 176–184.

Wyatt R (1982). Inflorescence architecture: how flower number, arrangement, and phenology affect pollination and fruit-set. American Journal of Botany 69: 585–594.

Xia Y, Xue B, Shi M, Zhan F, Wu D, Jing D, Wang S, Guo Q, Liang G, He Q (2020). Comparative transcriptome analysis of flower bud transition and functional characterization of EjAGL17 involved in regulating floral initiation in loquat. PLoS One 15(10): e0239382.

Yang T, Wang L, Li C, Liu Y, Zhu S, Qi Y, Liu X, Lin Q, Luan S, Yu F (2015). Receptor protein kinase FERONIA controls leaf starch accumulation by interacting with glyceraldehyde-3-phosphate dehydrogenase. Biochemical and Biophysical Research Communications 465(1): 77–82.

Yapo BM (2011). Rhamnogalacturonan-I: A structurally puzzling and functionally versatile polysaccharide from plant cell walls and mucilages. Polymer Reviews 51(4): 391–413.

Yu H, Ito Y, Zhao Y, Peng J, Kumar P, Meyerowitz E (2004). Floral homeotic genes are targets of gibberellin signaling in flower development. Biological Sciences 101(20): 7827–7832.

Zhang D, Ren L, Yue JH, Wang l, Zhuo LH, Shen XH (2013). A comprehensive analysis of flowering transition in *Agapanthus praecox* ssp. *orientalis* (Leighton) Leighton by using transcriptomic and proteomic techniques. Journal of Proteomics 80: 1–125.

Zu Y, Klasfeld S, Jeong CW, Jin R, Goto K, Yamaguchi N, Wagner D (2020). Terminal flower 1-FD complex target genes and competition with FLOWERING LOCUS T. Nature communications 11: 5118.

