## Supplementary figures and images for "Molecular mechanisms underlying the early steps of floral initiation in seasonal flowering genotypes of cultivated strawberry"

### Supplemental Figure S1

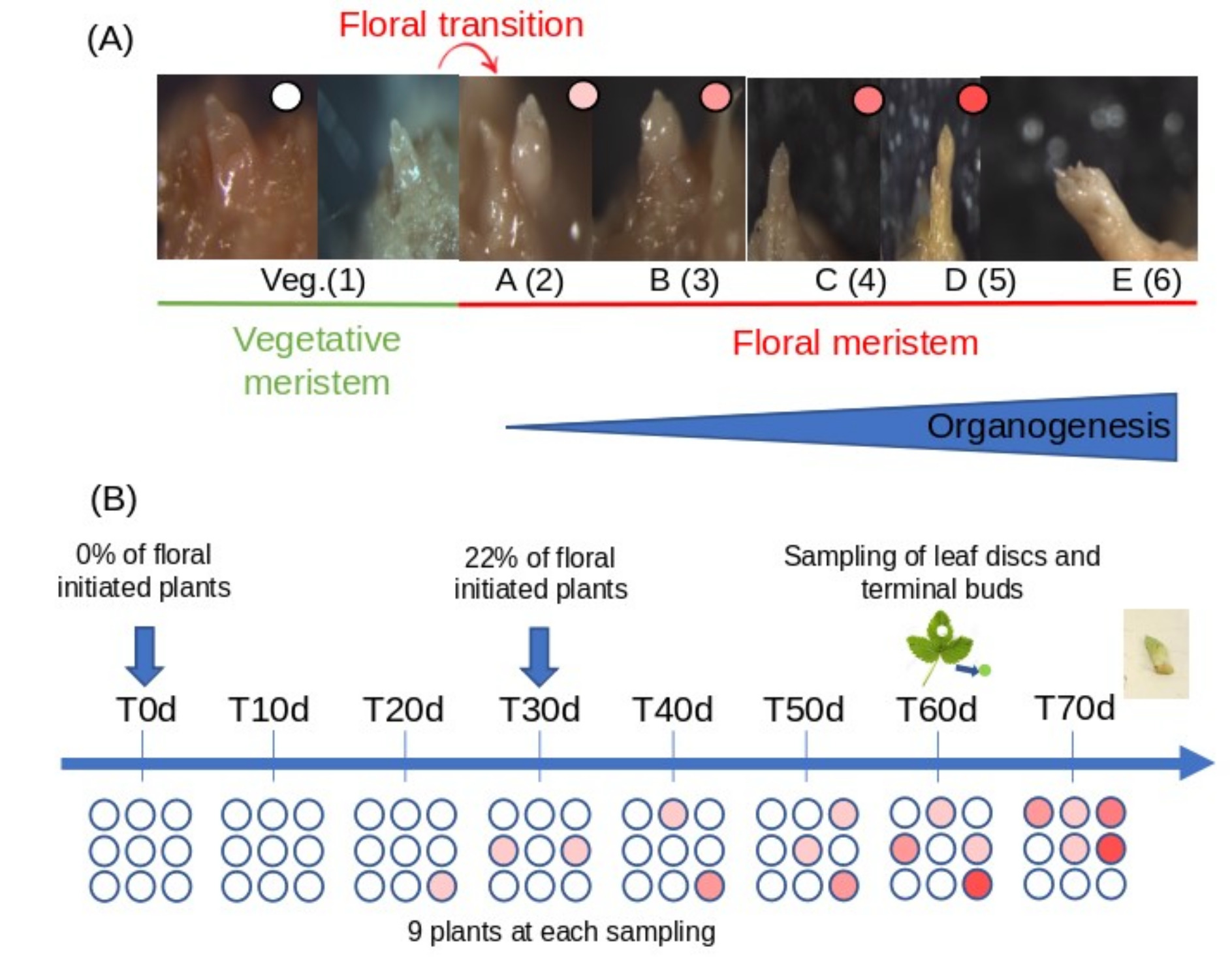

### Supplemental Figure S2

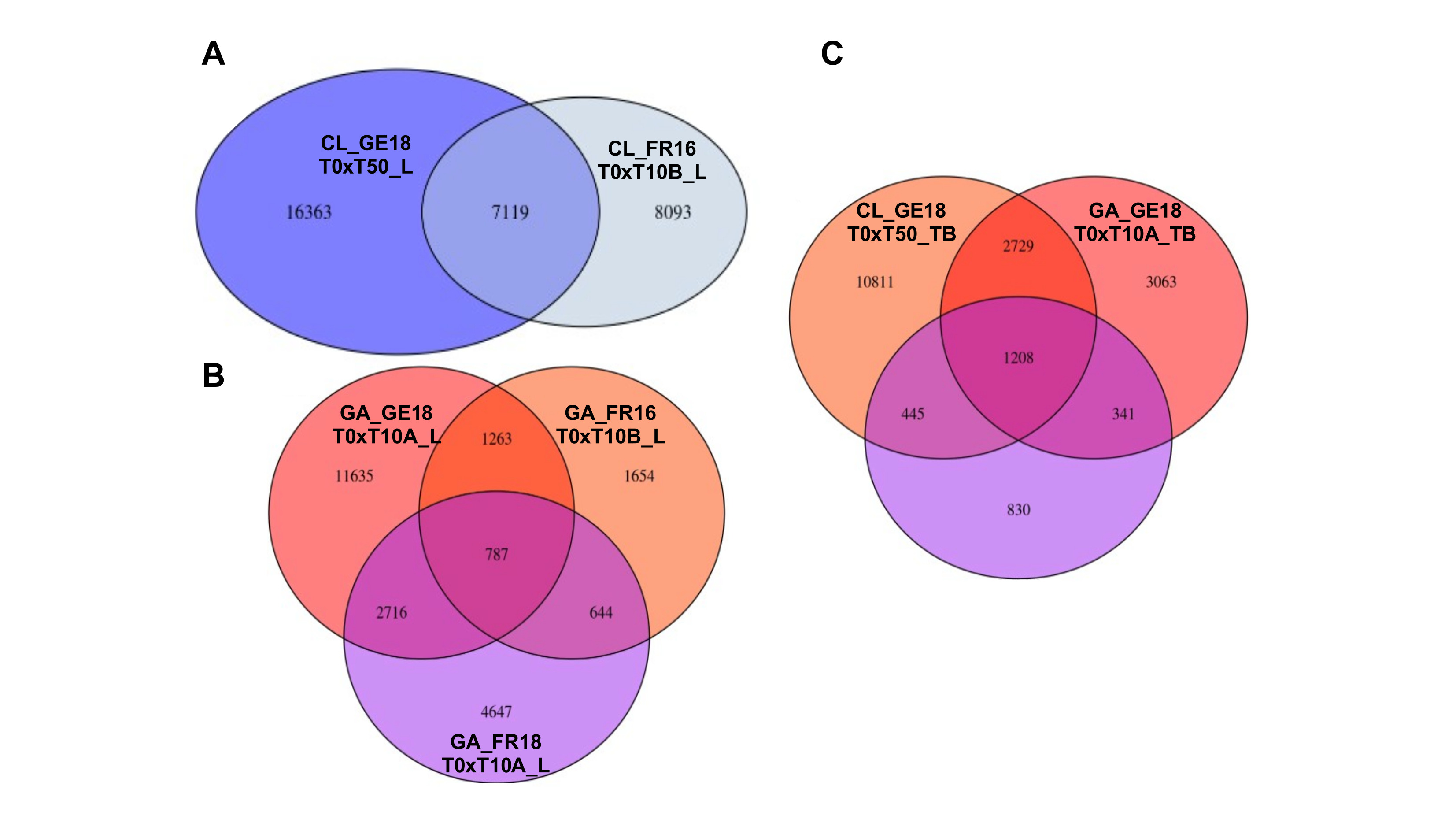

### Supplemental Figure S3

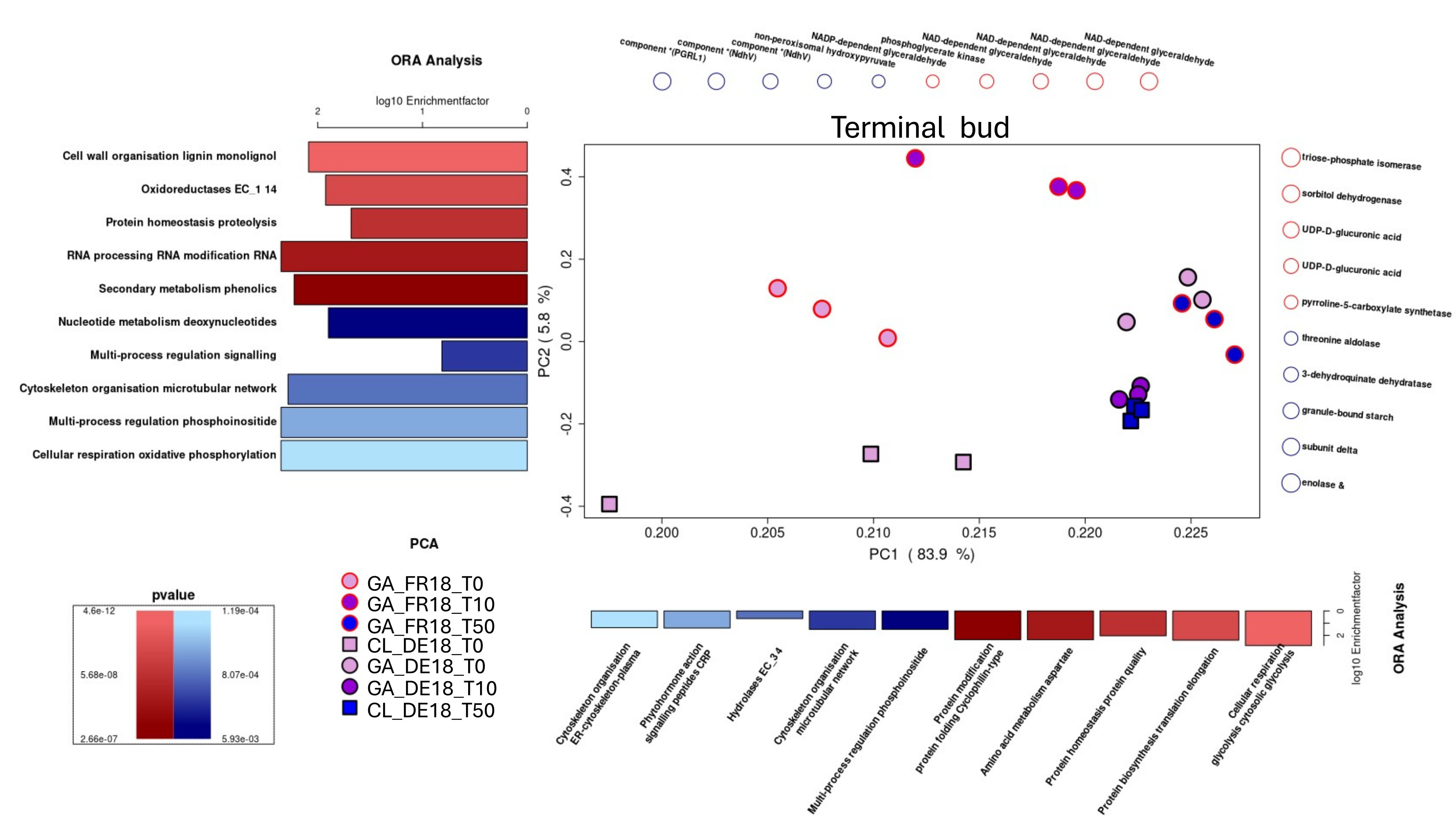
